# B cells are addicted to immunoglobulin production in the ER

**DOI:** 10.1101/2022.03.11.483940

**Authors:** Marieta Caganova, Hakan Taskiran, Yue Zong, Andreas Zach, Eleni Kabrani, Kristin Schmidt, Paul Goffing, Julia Jellusova, Klaus Rajewsky

## Abstract

Resting B cell dependence on the B cell antigen receptor (BCR) has been attributed solely to its signaling competence. We have recently shown that the presence of the BCR is vital for endoplasmic reticulum (ER) homeostasis and mitochondrial function both in primary B cells and Burkitt lymphoma (BL) cell lines. Unexpectedly, in BL Ramos cells this role has been shown to be independent of BCR signals from the cell membrane. Now, we provide evidence that also in mouse resting B cells ER-resident immunoglobulin (Ig) heavy chains control ER homeostasis, calcium storage and mitochondrial homeostasis through ER-mitochondria contacts. These findings demonstrate that BCR expression shapes cellular fitness already at the stage of assembly in the ER and uncover a new layer of B cell dependence on the production of Ig heavy chains in the ER. Sensing protein expression in the ER, with counterselection of compromised cells, might well operate also in other differentiated cells.

## Introduction

The BCR is composed of Ig heavy and light chains that assemble with Ig alpha (Igα) and beta (Igβ) in the ER to be subsequently transported to the cell membrane. Signals derived from the BCR are necessary for B cell maturation and egress from the bone marrow during B cell development (Rajewsky, 1996). The loss of such signals leads to a complete arrest of B cell development.

The evidence that BCR expression is essential also for the survival of mature resting B cells came from a series of conditional gene targeting experiments. In 1997, Lam et al. showed that the deletion of a pre-rearranged VDJ element (B1-8) and thus ablation of the BCR in mature B cells led to their disappearance due to apoptosis (Lam *et al*, 1997). This study uncovered a fundamental quality control step which ensures that the population of mature B cells is only composed of BCR expressing cells capable of responding to pathogens, and that non-functional B cells are eliminated.

However, the question whether it is a BCR-derived signal or the physical presence of the BCR that is vital for B cells remained unanswered. Subsequently Kraus et al. provided evidence that a BCR-derived signal is essential for B cell survival. Similar to conditional B1-8 deletion, the deletion or mutation of Igα (the signaling subunit of the BCR) led to a prompt disappearance of the targeted B cells from the periphery (Kraus *et al*, 2004).

Finally, Srinivasan et al identified the nature of the BCR maintenance signal in resting mature B cells in an elegant rescue experiment. The authors hypothesized that one of the well-known signaling pathways (PI3K, MAPK, NFκB), normally activated upon BCR activation through cognate antigen, might also mediate the maintenance signal from the BCR in resting B cells. Their results showed that concomitant BCR ablation and PI3K pathway activation, exclusively, led to a complete rescue of the B1-8 deleted B cells (Srinivasan *et al*, 2009).

Experiments using the Epstein Barr virus encoded LMP2A protein, able to mimic BCR-mediated signals, demonstrated that a BCR-like signal originating from a BCR unrelated molecule can support the development of mature resting, BCR-negative B cells in a mouse model (Caldwell *et al*, 1998). The level of LMP2A expression is, however, critical for B cell lineage decisions. Low LMP2A expression levels (from a D_**H**_ promoter, D_**H**_LMP2A) led to the development of B2-like BCR-negative B cells while high LMP2A expression (from a V_**H**_ promoter, V_**H**_LMP2A) directed B cells into a B1-like B cell fate (Casola *et al*, 2004).

Together, these findings supported the concept of a membrane BCR-mediated maintenance signal in mature resting B cells, termed “tonic signal”.

These experiments were, however, not designed to study the impact of the BCR deletion on the intracellular environment, specifically its impact on the endoplasmic reticulum (ER). A resting mature B cell carries 130.000 -250.000 BCRs on its surface (Yang & Reth, 2010; Treanor *et al*, 2010), with a turnover rate of approximately 20 h (Melchers, 1978). Thus, the maintenance of BCR expression on B cells depends on de novo BCR production in the ER. How such BCR production shapes the ER of resting cells and whether the loss of BCR has any impact on ER homeostasis and B cell fitness has long remained unaddressed.

Recently, we have shown that the presence of Ig heavy chain can be sensed in the ER, and that its expression correlates with ER homeostasis and mitochondrial function both in primary B cells and Burkitt lymphoma cell lines (Jumaa *et al*, 2020). Surprisingly, in the Burkitt lymphoma Ramos cell line this role was completely independent of BCR signals from the cell membrane (Jumaa *et al*, 2020). This prompted us to further investigate the role of ER-resident Ig heavy chains in primary resting B cells. In comparison to lymphoma cells, naïve B cells display low levels of metabolic activity and a small ER. Thus, the goal of the present study was to establish whether the presence of the Ig heavy chains shapes ER homeostasis in resting B cells, thereby contributing to cellular fitness and maintenance of a functional B cell pool. For this purpose, we used CRISPR-Cas9-mediated mutagenesis of BCR components in ex vivo purified mature resting B cells and an in vivo mouse model which enabled us to distinguish between cell membrane-bound and the ER-resident functions of Ig heavy chains. We provide evidence that in resting B cells Ig heavy chains are sensed in the ER, control ER homeostasis, ER-mitochondria contact sites and support mitochondrial function.

## Results

### ER-resident Ig heavy chains provide competitive advantage to B cells

To understand if Ig heavy chains from within the ER can provide pro-survival signals to naïve splenic B cells, we performed CRISPR-Cas9 mutagenesis using Cas9 protein with BCR specific sgRNAs. We used resting primary B cells isolated from B1-8i transgenic mice that carry a pre-rearranged VDJ element in its physiological position in the IgH locus (B1-8). We targeted the VDJ region (sgB1-8) or CD79b (sgIgβ) aiming to preclude expression of Ig heavy chains or Igβ, a BCR component involved in BCR membrane anchorage and signal transduction, respectively (**Fig 1a**: scheme). Successful targeting with sgB1-8 or sgIgβ led to BCR loss at the cell surface. While Igβ targeted B cells retained intracellular Ig heavy chains, B cells targeted with sgB1-8 were completely devoid of Ig heavy chains, as both IgM and IgD were lost upon successful targeting (**Fig 1b, SFig 1a**). Furthermore, functional loss of surface IgM was demonstrated by the inability of sgB1-8 and sgIgβ targeted B cells to induce Syk phosphorylation upon anti-IgM stimulation (**Fig 1c)**. Surface BCR-deleted B cells with retained intracellular Ig heavy chains (sgIgβ) showed significantly better competitive fitness over 6 days of culture with BAFF as compared to sgB1-8 (**Fig 1d**), suggesting that intracellular Ig heavy chains provide a pro-survival signal. Similar results were obtained using the Burkitt lymphoma cell line DG75 expressing the Cas9 protein. DG75 cells were transduced with sgRNA expressing vectors targeting the constant region CH1 (sgCH1) or transmembrane region M1 (sgM1) of human IgM (**Fig S1b)** which led to the loss of surface IgM in both cases, while intracellularly, ER-resident Ig heavy chains were absent (sgCH1) or retained (sgM1), respectively (**Fig S1c**). Functionally, surface IgM deletion was evident from the inability of successfully targeted DG75 cells to respond to anti-IgM stimulation by Syk phosphorylation (**FigS1d**). The retention of ER-resident Ig heavy chains in surface IgM ablated DG75 cells (sgM1) led to a significantly improved fitness as compared to the complete IgM-ko cells (sgCH1) (**FigS1e**). To test if ER-resident Ig heavy chains impact mitochondrial function we measured oxygen consumption rates of the targeted and control DG75 cells and observed that ER-resident Ig heavy chains (sgM1) are sufficient to support normal mitochondrial metabolism of these cells (**Fig S1f**). We conclude that intracellular Ig heavy chains provide a fitness advantage to surface BCR deleted B cells.

**Figure 1.**
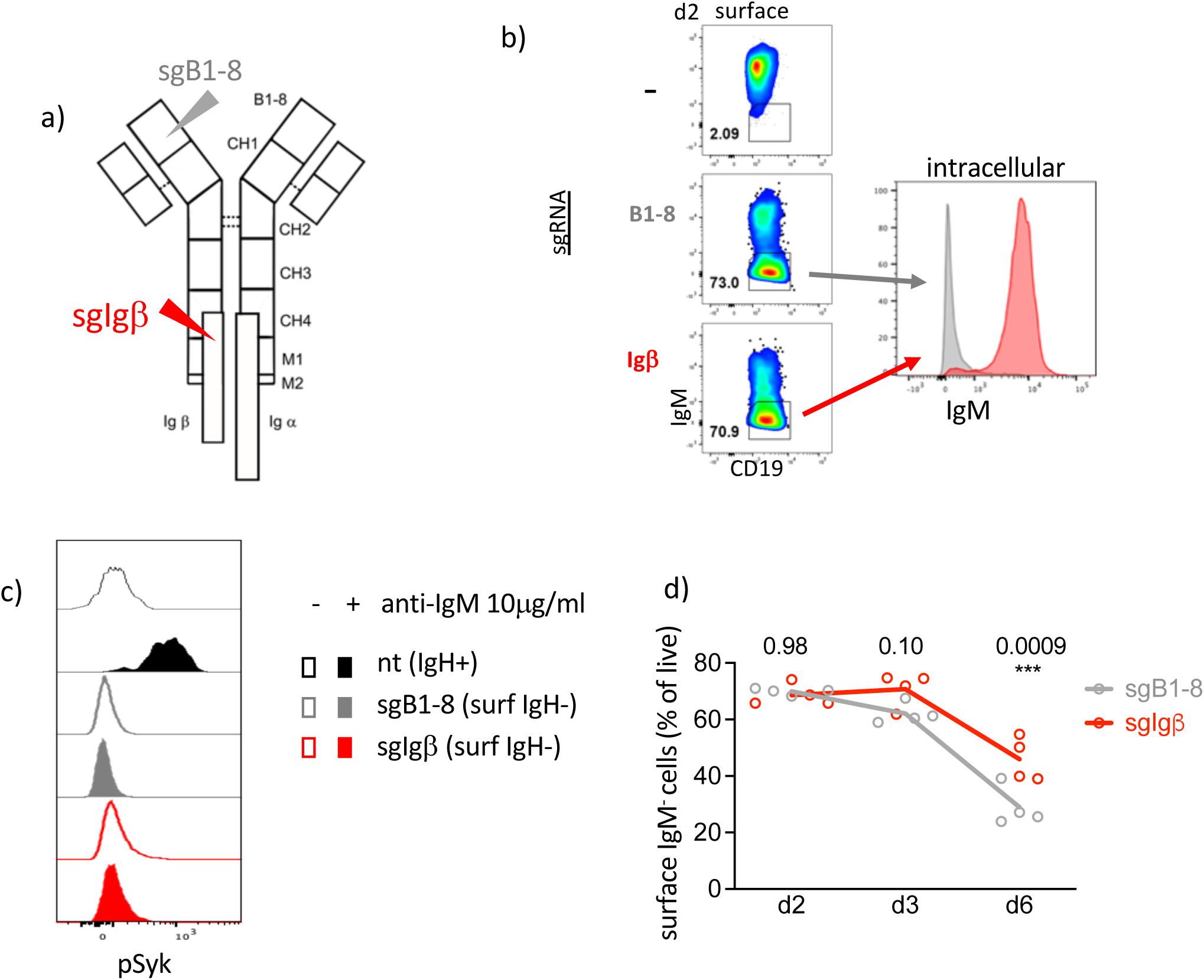
Intracellular Ig heavy chains provide competitive advantage to B cells. (a) Schematic depiction of BCR targeting strategy with sgRNAs specific for B1-8 (grey arrowhead) or Igβ (red arrowhead). (b) A representative flow cytometry analysis of mature resting B cells stained with anti-CD19 and anti-IgM on surface and intracellularly on d2 upon mock, sgB1-8 or sgIgβ targeting, cultured with BAFF. (c) Phospho-flow analysis of pSyk in mature resting B cells on d3 upon transduction using mock, sgB1-8 or sgIgβ with or without anti-IgM activation; cultured with BAFF. (d) Fitness of mature resting surface IgM-negative B cells targeted using sgB1-8 or sgIgβ and cultured with BAFF for 6 days. Results were tested for significance with ANOVA. Data obtained from 4 mice in 2 independent experiments.

### ER and metabolic defects in B cells lacking ER-resident Ig heavy chains

Previously, we have shown that mitochondrial metabolism and ER homeostasis is impaired upon Ig heavy chain loss and that this phenotype can be rescued by the re-expression of immunoglobulins in the absence of the BCR signaling subunits Igα and Igβ in the Ramos B cell line (Jumaa *et al*, 2020). However, whether intracellular Ig heavy chains can provide such rescue signals in primary B cells in vivo remained unknown. Primary B cells are fundamentally different to Ramos cells in that they harbor low ER mass and display little mitochondrial activity. To assess whether Ig heavy chain-dependent ER-homeostasis contributes to cellular fitness in resting B cells, we took advantage of the D_**H**_LMP2A mouse (Casola *et al*, 2004), in which the coding sequence of the EBV gene LMP2A is knocked-in into the IgH locus, replacing the D_**H**_ genes in homozygous fashion (D_**H**_LMP2A) and providing a “tonic”, BCR-like, survival signal that enables BCR-negative B cells to egress from the bone marrow and populate the spleen as mature B2-like B cells (**Fig 2a: scheme**). Despite their ability to overcome the block in B cell development upon IgH-inactivation, absolute splenic B cell numbers in D_**H**_LMP2A mice are significantly reduced as compared to controls, suggesting their decreased fitness (Casola *et al*, 2004; and **Fig 2b**). We tested ER homeostasis and mitochondrial metabolism of D_**H**_LMP2A B cells and observed that the defects of IgH-ko B cells (Jumaa *et al*, 2020) are reproduced in D_**H**_LMP2A B cells, namely, D_**H**_LMP2A B cells showed reduced ER tracker staining intensity (**Fig 2c**) and reduced mRNA expression of the ER-resident *Hspa5* (BIP) chaperone (**Fig 2d**). ER-dependent processes are known to shape mitochondrial respiration. To study mitochondrial function, we analyzed the basal and maximal oxygen consumption rate and mitochondria dependent ATP production in D_**H**_LMP2A B cells. Consistent with the hypothesis that an atrophied ER limits mitochondrial function we found D_**H**_LMP2A B cells display reduced mitochondrial activity similar to IgH-ko B cells (**Fig 2e,f**). Thus, LMP2A-driven pro-survival signals cannot fully rescue ER homeostasis and metabolic fitness upon BCR loss.

**Figure 2.**
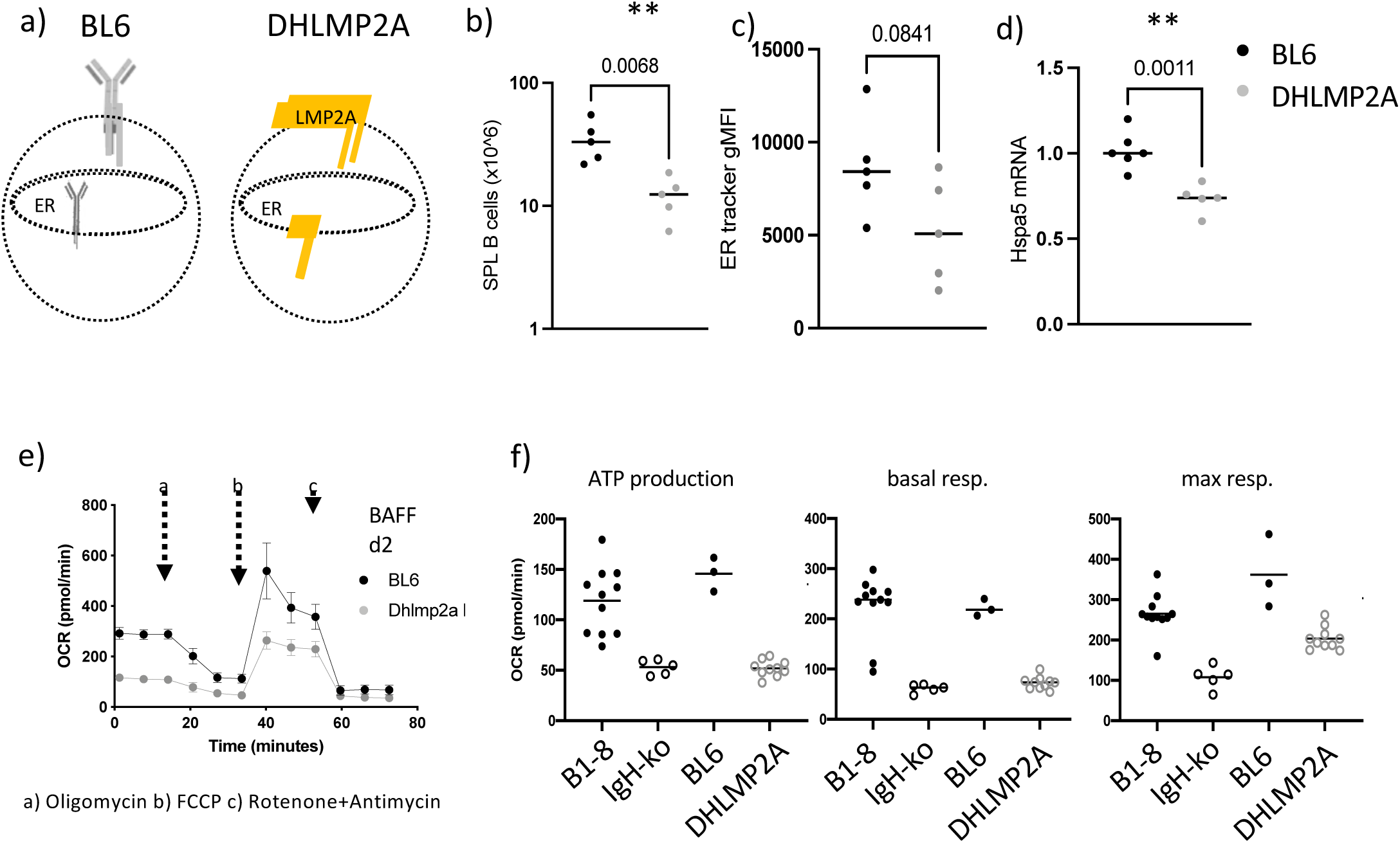
Lack of ER resident Ig heavy chains leads to defects in ER and metabolic homeostasis. (a) Schematic depiction of B cells from control (BL6) and D_**H**_LMP2A mice. (b) Total splenic B cells of BL6 (black dots) and D_**H**_LMP2A (grey dots) mice. Results were tested for significance with t-test. (c) ER tracker geometric mean fluorescence intensity of BL6 (black dots) and D_**H**_LMP2A (grey dots) splenic B cells. Results were tested for significance with Mann-Whitney test. (d) *Hspa5* (BIP) mRNA expression levels measure by qRT-PCR normalized with *Rplp0*. Results were tested for significance with t-test. (e) Oxygen consumption rate measured by Mito stress test Seahorse assay on B cells purified from BL6 and D_**H**_LMP2A B cells cultured in presence of BAFF for 2 days. (f) The calculations of ATP production, basal respiration and maximal respiration are detailed in Material and Methods section. Data obtained from a minimum of 3 mice in 2 independent experiments.

### Intracellular Ig heavy chains are sensed in the ER and improve B cell fitness

We hypothesized that the lack of ER-resident Ig heavy chains might be responsible for impaired ER and metabolic homeostasis of D_**H**_LMP2A B cells and that Ig heavy chain expression should rescue, at least partially, these defects. To test this hypothesis, we bred D_**H**_LMP2A, B1-8i and RAGko mice to obtain a compound mutant B1-8i/D_**H**_LMP2A; RAGko (BDR) in which D_**H**_LMP2A provides the tonic signal allowing B cell maturation; B1-8i assures Ig heavy chain expression and the RAGko background prohibits Ig light chain gene rearrangement. In this setting, D_**H**_LMP2A allows surface Ig-negative B cell maturation with ER-resident Ig heavy chains, unable to translocate to the cell surface (**Fig 3a:** scheme). As controls, D_**H**_LMP2A/D_**H**_LMP2A; RAGko (DDR) animals, which do not produce Ig heavy chains were used. As a wild type control, C57BL/6 (hereafter indicated as BL6) mice were used. We characterized the B cell subsets in the newly generated compound mutants and observed the surface phenotype of BM and SPL B cell subsets to be comparable among BL6, D_H_LMP2A, DDR and BDR mice (**Fig S2a,b**), with the expected mild increase of CD19 and decrease of B220 expression on the surface of D_**H**_LMP2A expressing B cells as reported previously (Casola *et al*, 2004). The high expression of CD21, CD23 and CD38 on the mature (CD19^+^AA4.1^-^) splenic B cells from BL6, D_**H**_LMP2A, DDR and BDR mice confirmed their B2-like cell phenotype (**Fig 3b,SFig 2b**. Virtually all mature splenic B cells from BDR mice expressed ER-retained Ig heavy chains unable to translocate to the B cell surface (**Fig 3b,SFig 2c,d**. Moreover, functionally, we confirmed the absence of surface IgM in BDR B cells by their inability to respond to anti-IgM stimulation by calcium efflux (**Fig 3c**). Next, we determined absolute splenic B cell numbers and observed that the expression of ER-retained Ig heavy chains led to a significant normalization of mature B cell numbers in BDR as compared to DDR mice (**Fig 3d**). To exclude that the pre-BCR plays a role in the rescue of BDR B cells we stained splenic cells from BL6, BDR and DDR mice with an anti-CD179a (VpreB) antibody intracellularly (given the notoriously low expression of pre-BCR on the cell surface (Brouns *et al*, 1996)). As positive controls we used bone marrow cells where we observed intracellular VpreB expression exclusively in progenitor B cells (**SFig 2e)**. VpreB was completely absent from splenic B cells from all, BL6, DDR and BDR mice (**Fig 2e**). Therefore, we can exclude that the pre-BCR plays any role in the improved mature B cell fitness in BDR mice. Hence, we provide evidence that ER-resident Ig heavy chains are sensed in the ER and improve the fitness of B cells in this model.

**Figure 3.**
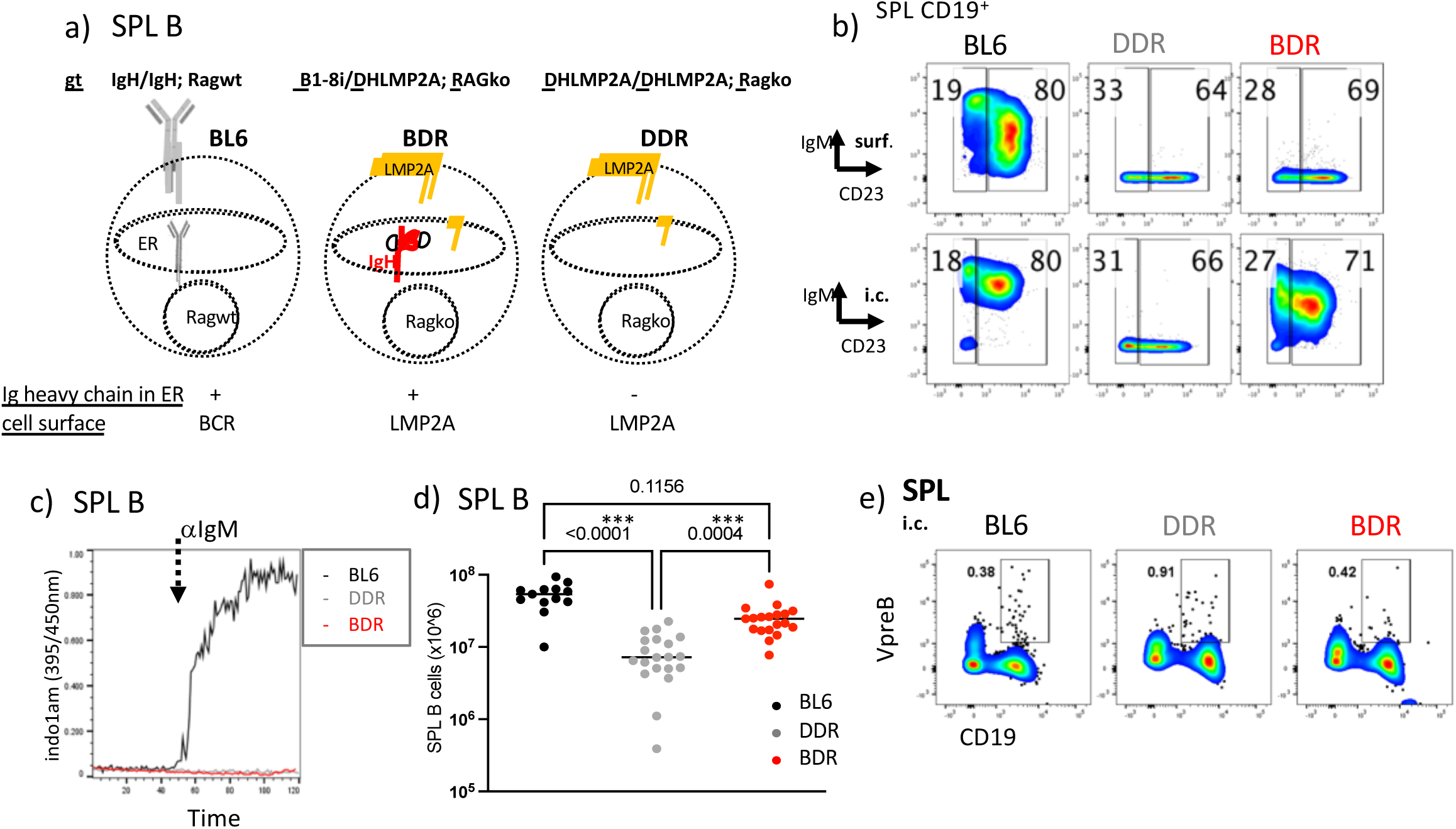
Intracellular Ig heavy chains are sensed in the ER and improve B cell fitness. (a) Schematic depiction of B cells from BL6, BDR and DDR mice. (b) Representative flow cytometry analysis of splenic B cells from BL6, BDR and DDR mice stained with anti-IgM and anti-CD23 on surface (upper panel) and intracellularly (lower panel). (c) Representative flow cytometry analysis of calcium efflux of BL6, BDR and DDR B cells upon anti-IgM stimulation. (d) Total splenic B cells of BL6 (black dots), DDR (grey dots) and BDR (red dots) mice. Results were tested for significance with Kruskal-Wallis test. (e) Representative flow cytometry analysis of splenic cells from BL6, BDR and DDR mice stained with anti-CD19 and anti-CD179a (VpreB) intracellularly. Data obtained from a minimum of 3 mice in 2 independent experiments.

### ER-resident Ig heavy chains sustain ER homeostasis and mitochondrial metabolism

We have shown that B cells lacking ER resident Ig heavy chains display defects in ER homeostasis and mitochondrial function. Next, we decided to test whether ER-resident Ig heavy chains can rescue these defects. We stained BL6, DDR and BDR B cells with ER-tracker and confirmed a significantly reduced ER mass in DDR B cells as compared to controls. In BDR B cells we observed increased ER tracker signal intensity as compared to DDR B cells, suggesting a partial rescue of ER mass, although this rescue did not reach statistical significance (**Fig 4a**).

**Figure 4.**
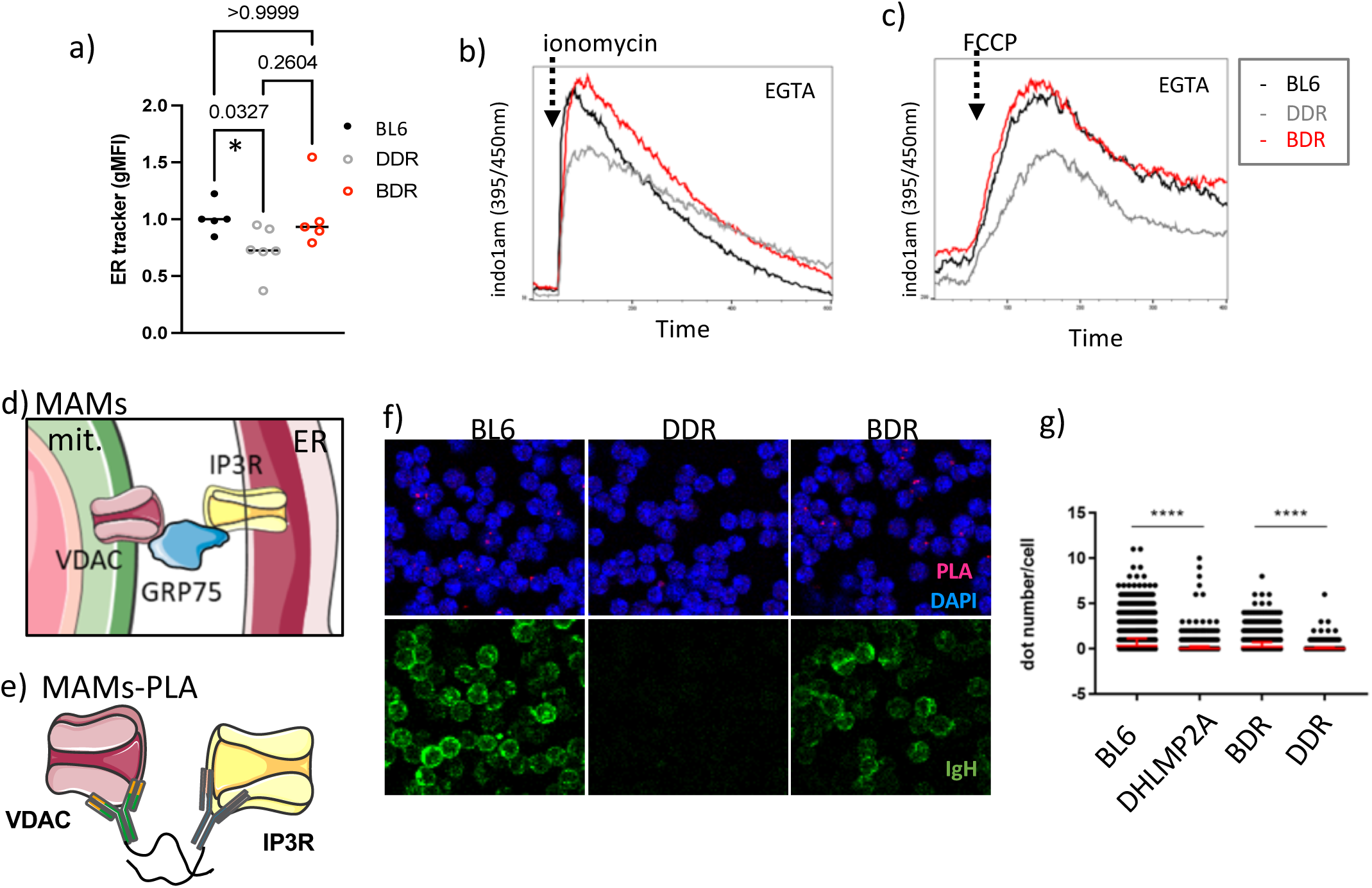
ER-resident Ig heavy chains improve ER homeostasis and MAMs. (a) ER tracker geometric mean fluorescence intensity of BL6, DDR and BDR splenic B cells. Results were tested for significance with Kruskal-Wallis test. (b) Representative flow cytometry analysis of calcium efflux upon ionomycin stimulation of control, DDR and BDR B cells in presence of EGTA. (c) Representative flow cytometry analysis of calcium efflux upon FCCP stimulation of control, DDR and BDR B cells in presence of EGTA. (d) Schematic depiction of mitochondria-associated membranes (MAM). (e) Schematic depiction of proximity ligation assay (PLA) to detect MAMs. (f) Representative fluorescent microscopy images showing B cells upon MAM-PLA assay (PLA signal in red), stained with DAPI (blue) in upper panel; and anti-IgM (green) in lower panel. (f) Quantification of PLA signals from (e) was performed with CellProfiler Software. Results were tested for significance with ANOVA. Data obtained from a minimum of 3 mice in 3 independent experiments.

The ER is the most prominent calcium storage organelle in non-muscle cells. We asked if the impaired ER homeostasis caused by the lack of Ig heavy chains has an impact on calcium storage capacity in the ER. To test the calcium storage capacity in ER-resident Ig heavy chain proficient and deficient B cells, we stained BL6, DDR and BDR B cells with the calcium indicator indo1am and induced calcium efflux by ionomycin treatment in presence of the extracellular calcium chelator EGTA to prevent calcium influx from the extracellular space. We observed reduced calcium efflux in DDR B cells, indicating their lower calcium storage capacity as compared to controls (**Fig 4b**). This reduction was readily rescued in BDR B cells (**Fig 4b**); thus, the ER calcium storage capacity is controlled by ER-resident Ig heavy chains. The ER is an essential calcium provider for the mitochondria, which depend on ER-mediated calcium supply for proper tricarboxylic acid cycle activity. To test if mitochondrial calcium levels are affected by the ER calcium storage capacity, we stained BL6, DDR and BDR B cells with indo1am and induced calcium efflux from the mitochondria by disrupting their membrane potential using Carbonyl cyanide-p-trifluoromethoxyphenylhydrazone (FCCP) in the presence of EGTA. We observed reduced mitochondrial calcium efflux in DDR B cells as compared to BL6, while the levels were normalized in BDR B cells (**Fig 4c**). Importantly, the reduced ER and mitochondrial calcium efflux was also seen in primary B cells upon in vitro TAT-cre induced IgH ablation (**SFig 3a,b**); as controls B cells from R26-hCD2^flSTOP^ mice with induced hCD2 surface expression upon TAT-cre treatment were used.

ER and mitochondria form well-defined contact sites, also termed mitochondria-associated membranes (MAMs), which are essential for communication and calcium flow between the ER and mitochondria (**Fig 4d scheme**). Given the correlation of reduced calcium levels between ER and mitochondria in B cells lacking ER-resident Ig heavy chain, we decided to test if ER-resident Ig heavy chains affect MAM formation. To quantify MAMs, we performed a proximity ligation assay (PLA) using antibodies specific for proteins involved in MAMs (Bantug *et al*, 2018); IP3R (ER membrane) and VDAC (mitochondrial membrane) in BL6, DDR and BDR purified B cells (**Fig 4e scheme**). We observed significantly decreased numbers of MAMs in the absence of ER-resident Ig heavy chains (in DDR B cells) (**Fig 4f,g**). To exclude that the impaired MAM formation is due to reduced levels of the IP3R and VDAC proteins in DDR B cells, we performed immunoblotting experiments and verified that the expression of the tested proteins was comparable between BL6 and DDR B cells (**SFig 3c,d**). Next, we observed a partial, yet significant, rescue of MAMs upon ER-resident Ig heavy chain expression in BDR as compared to DDR B cells (**Fig 4f,g**). Post-PLA, the cells were stained with anti-IgM antibody to confirm the absence or presence of intracellular Ig heavy chains in DDR and BDR B cells, respectively. Taken together, we show that the expression of ER-resident Ig heavy chains is required in primary naïve B cells for ER homeostasis, its calcium storage capacity, contacts to mitochondria and finally, correct mitochondrial calcium levels.

In conclusion, we provide evidence that B cells depend on ER-resident Ig heavy chain expression for proper ER and mitochondrial homeostasis.

## Discussion

The notion that a tonic signal, i.e. a maintenance signal originating from the membrane-bound BCR, is necessary for mature resting B cells to persist in the periphery has strongly increased our understanding of how a pool of healthy B cells is maintained. B cells unable to form a functional BCR lose their ability to respond to antigen and need to be eliminated. However, the impact of BCR loss on ER-associated processes has not been studied.

Recently, we provided first evidence that immunoglobulin expression in the ER rescued impaired ER homeostasis and mitochondrial metabolism in BCR-ablated Ramos cells, and that this rescue was independent of membrane BCR-mediated signals (Jumaa *et al*, 2020). However, evidence that Ig heavy chain expression within the ER functionally affects primary B cells was still missing.

To address the ER-associated function of Ig heavy chains, here, we employed three independent experimental systems: 1) CRISPR-Cas9 mediated mutagenesis in ex vivo purified resting mature B cells; 2) a mouse model in which B cells are maintained through a BCR-like signal independent of Ig expression and 3) CRISPR-Cas9 mediated mutagenesis in the Burkitt lymphoma cell line DG75. We provide evidence that Ig heavy chains are sensed from within the ER and i) provide competitive advantage to B cells, ii) maintain ER and calcium homeostasis, iii) control ER-mitochondria contacts and iv) improve mitochondrial metabolism in B cells. This extends the conclusions of the Jumaa *et al*, 2020 study to mature resting B cells.

How do our results go in line with previously published data? Kraus et al provided first evidence that signals originating from the membrane-bound BCR are required for mature B cell survival. The authors compared the effect of complete Ig heavy chain (B1-8) or Igα (BCR signaling unit) deletion and in both cases B cell survival was strongly reduced with an estimated half-life of Igα-deleted B cells between 3-6 days (Kraus *et al*, 2004). Based on our results (Fig 1d) we speculate, that the half-life of B1-8 deleted B cells might be even shorter than that of Igα-deleted B cells. There is however, in general, no discordance here, as also our data indicate that although ER-resident Ig heavy chains are able to improve B cell fitness over a short period of time, they are insufficient to provide a full survival stimulus to B cells in the absence of a signal originating from the membrane-bound BCR (Fig 1d).

Srinivasan et al. showed that survival and persistence of BCR-deleted B cells was rescued completely by PI3K signals (Srinivasan *et al*, 2009). It is important to note that activation of PI3K is sufficient to upregulate BIP mRNA expression in B cells in vitro (Jumaa *et al*, 2020). This indicates, that PI3K may act on two, probably independent levels: i) mimicking the BCR-derived tonic survival signal and ii) inducing the expression of selected ER-resident proteins to maintain ER homeostasis in resting B cells. Its dual function could permit activated PI3K to provide a complete rescue to BCR-less B cells.

How is ER homeostasis controlled by ER-resident Ig heavy chains? During BCR assembly in the ER, the CH1 region of the Ig heavy chain molecule is bound by two BIP chaperon proteins until folded Ig light chains finally release the Ig heavy chain from BIP (Lee *et al*, 1999; Haas & Wabl, 1983) and the fully assembled BCR translocates through the Golgi apparatus onto the cell membrane (Hombach *et al*, 1990). BIP is one of the central players in the unfolded protein response pathway which is activated upon increased ER-stress (reviewed in Walter & Ron, 2011). BIP’s physiological levels, vital for proper ER homeostasis, are strictly regulated by feedback mechanisms (Gülow *et al*, 2002; Bakunts *et al*, 2017). We hypothesize that the Ig heavy chain-BIP equilibrium plays a major role in B cell ER homeostasis. Upon sudden Ig heavy chain loss, this equilibrium is disrupted, and the resulting excess of free BIP protein might activate endogenous feedback mechanisms that promptly decrease de novo BIP protein synthesis (we observed decreased BIP expression in B cells lacking ER-resident Ig heavy chains (Fig. 2d; and Jumaa *et al*, 2020). We speculate, that the reduction of BIP protein might result in reduced ER mass and calcium storage capacity in B cells lacking ER-resident Ig heavy chains.

Moreover, BIP protein has been shown to bind, and improve the stability of, ER-mitochondria contact sites (MAMs) (Hayashi & Su, 2007), which were significantly reduced in the absence of ER-resident Ig heavy chains (DDR) and readily rescued by Ig heavy chain expression in the ER (BDR) (Fig 4f, g). The ER as the most important calcium storage organelle in non-muscle cells (Koch, 1990) is, through the MAMs, responsible for the flow of calcium into the mitochondria (reviewed in Hajnóczky *et al*, 2002). The reduced mitochondrial calcium levels observed in B cells lacking ER-resident Ig heavy chains might be therefore explained by the reduction of MAMs and/or reduced calcium storage capacity of the ER.

It is important to note here, that some defects observed in B cells lacking ER-resident Ig heavy chains (DDR) were only partially rescued upon the expression of ER-resident Ig heavy chains in BDR B cells (Fig 3d, 4a, 4f). We suspect that the lack of Ig light chains in BDR B cells might be responsible for such incomplete rescue. Further experiments are necessary to address the role of Ig light chains in ER homeostasis.

Taken together, we conclude that B cells, beyond the maintenance signal derived from the membrane-bound BCR, depend on immunoglobulin production in the ER. From the early stages of B cell development when the first Ig gene rearrangements are completed, B cells turn on immunoglobulin production that tunes their ER homeostasis. A sudden loss of Ig heavy chains in mature B cells is sensed in the ER and leads to an impairment of the ER machinery and cellular metabolism. Our study, thus, uncovers a state of addiction of B cells to the production of Ig heavy chains in the ER.

Our findings that BCR-dependent ER homeostasis is central to B cell fitness in both resting mature and transformed B cells has implications for various aspects of B cell biology. First, in B cell physiology, the processes of B cell maturation, activation and differentiation which lead to substantial changes of the ER-protein load, may directly impact the metabolic activity and flexibility of these cells. The newly discovered role of Ig heavy chains in shaping ER homeostasis and ultimately mitochondrial function, adds to our understanding of how metabolism is controlled during different stages of B cell development and differentiation. Second, in B cell pathology, the newly discovered molecular players in the control of mitochondrial calcium homeostasis might represent therapeutic targets for the treatment of B cell malignancies resistant to inhibitors targeting classical BCR-dependent signaling pathways.

Finally, the present work raises the question whether the principle of ER-mediated addiction to, and thus monitoring of, the production of specialized proteins, in concert with counterselection of compromised cells, extends to cell types beyond antibody producing cells.

## Materials and Methods

### Mouse lines

D_**H**_LMP2A (Casola *et al*, 2004), RAG2-ko (Hao & Rajewsky, 2001), B1-8i (Sonoda *et al*, 1997), B1-8^f^ (Lam *et al*, 1997); EμBcl2 mice (Strasser *et al*, 1991), R26-hCD2^flSTOP^ (Calado *et al*, 2012) mice were used. Animals were used, bred and kept in accordance with the German Animal Welfare Act, approved by the regional council (LAGeSO n. G0196/19 and n. X 9002/20).

### Isolation of primary resting B cells and in vitro treatments

Resting splenic B cells were purified by negative selection using autoMACS proSeparator or LS magnetic columns (Miltenyi) with CD43 depletion beads (Miltenyi) or using EasySep™ Mouse B Cell Isolation Kit (StemCell) with EasyEights™ EasySep™ magnet (StemCell). Primary B cells were plated at 2-3×10^6^ cells/ml in primary B cell medium (DMEM (Gibco) containing 10% FBS (Sigma), 2 mM Glutamax, 1 mM Sodium Pyruvate, 1 mM HEPES (Gibco), 1x NEAA (Gibco), 52 μM beta-mercaptoethanol (Sigma), 10 μg/ml Gentamicin (Lonza) and 100 ng/ml BAFF (Peprotech)).

TAT-cre treatment was used to induce deletion of pre-arranged VDJ (B1-8) in B1-8^f^; EuBcl2 or activation of hCD2 expression in R26hCD2^flSTOP^ B cells. Purified B cells were washed in PBS and transduced with 50 μg/mL TAT-Cre at a density of 5 × 10^6^/ml for 45 min at 37 °C in serum-free, animal-defined component-free medium (HyClone), prewarmed to 37 °C. CRISPR-Cas9 mediated mutagenesis in primary B cells was performed electroporating 10^6^ B cells with 50pmol Cas9 protein (homemade) with 100pmol sgRNA (sgB1-8 AGGATTGATCCTAATAGTGG, sgIgβ AGGTCACTGCTTGTCATTGC, IDT) in nucleofection buffer (5 mM KCl, 10 mM MgCl_2_; 20 mM HEPES; 90 mM Na_2_HPO_4_/NaH_2_PO_4_ pH7.2; 24 mM Sodium Succinate) using Amaxa™ 4D-Nucleofector™(Lonza). Electroporated B cells were culture in 200ul primary B cell medium for 6 days.

### CRISPR/Cas9 in DG75

CRISPR/Cas9 mediated knockout of IgM in DG75 cells was performed as described previously (Jumaa *et al*, 2020). To target the M1 region sgRNA_hIgHM_M1 (GAGGTGAGCGCCGACGAGGA) was used.

### Flow cytometry

Single cell suspensions were blocked with TruStain fcX (93), stained with fluorescently conjugated antibodies for 15 min in FACS buffer (PBS+1%BSA+0.09%NaN3), washed twice and acquired on BD Fortessa. Data were analyzed using FlowJo software. Primary mouse B cells were stained with anti-B220 (RA3-6B2), anti-CD19 (6D5), anti-IgM (II/41), Fab fragment anti-IgM (Jackson), anti-IgD (11-26), anti-CD23 (B3B4), anti-CD179a (R3), anti-CD21 (7E9), anti-CD43 (S7), anti-AA4.1 (AA4.1), anti-CD38 (37.51). Human DG75 cells were blocked with Human TruStain fcX and stained with anti-IgM (MHM-88). For intracellular staining BD Cytofix/Cytoperm™ (BD Biosciences) was used according to manufacturer’s protocol. For phospho-flow, primary resting B cells were stimulated with 10 μg/ml AffiniPure F(ab’)_2_ Fragment Goat Anti-Mouse IgM (JacksonImmuno), DG75 cells with 5 μg/ml anti-human IgM (DIANOVA) for 3 min. BD Cytofix/Cytoperm™ (BD Biosciences) was used and samples were stained with anti-pSyk (17A/P_ZAP70) and anti-Syk (5F5) in the presence of TruStain fcX (93) and Protease/Phosphatase inhibitor (CST).

### Immunoblot

5×10^6^ primary cells were lysed in 100μl RIPA buffer (50mM TRIS pH 8.0, 150mM NaCl, 1% Triton-X, 0.5% sodium deoxycholate, 0.1% SDS) containing Protease/Phosphatase inhibitor (CST) and ultrasonicated, and debris was removed by centrifugation. Supernatant was mixed with equal volume of 2X SDS buffer (20% glycerol, 4% SDS, 10% beta-mercaptoethanol, 0.01% bromophenol blue in 100 mM Tris-HCl, pH 6.8) and incubated at 99 °C for 10 min, then loaded to SDS-PAGE gel and transferred to PVDF membranes using Trans-Blot® Turbo™ Transfer System (Biorad). Primary antibodies anti-VDAC (D73D12, 1/1000) and anti-IP3R1 (E-8, 1/1000) were incubated in 5% BSA in TBS + 0.1% Tween (TBST) overnight at 4 °C. HRP-conjugated secondary antibody was diluted in 5% milk in TBST and incubated for 1 hour at room temperature. For signal detection SuperSignal West Femto (Thermo Scientific) was used.

### qRT-PCR

Total RNA was extracted using Mini RNeasy Kit (QIAGEN GmbH) with on-column DNA-se digestion and cDNA was synthetized using SuperScript IV (Thermo Scientific). 10-50ng cDNA were used for amplification with Fast SYBR™ Green Master Mix on StepOne Plus (Applied Biosystems) using specific primers (Rplp0_F TTCATTGTGGGAGCAGAC, Rplp0_R CAGCAGTTTCTCCAGAGC, Hspa5_F TTCAGCCAATTATCAGCAAACTCT,Hspa5_R TTTTCTGATGTATCCTCTTCACCAGT). ΔΔCT method was used to quantify mRNA expression.

### Proximity Ligation Assay (PLA)

Anti-VDAC1 (20B12AF2) (Abcam) and Anti-IP3R1 (E-8) (Santa Cruz Biotechnology) antibodies were used to generate PLA probes using Duolink in Situ Probemaker (Sigma). B cells were settled on PTFE diagnostic slides (Thermo Fisher Scientific) for 30 min at 37°C, fixed with 4% paraformaldehyde (Thermo Fisher Scientific) in PBS for 15 min, permeabilized with 0.5% saponin (*Quillaja* bark, Sigma) in PBS for 30 min and blocked with Duolink Blocking Solution (Sigma) for 1 hour. Cells were incubated with Anti-VDAC1 (1/100) and Anti-IP3R1 (1/100) PLA probes overnight at 4°C. PLA was performed according to the manufacturer’s instructions (Duolink, Sigma). Cells were then incubated with anti-IgM antibody (1/100) (Jackson Immuno Research) for 30 min at 4°C and mounted in Fluoromount-G with DAPI (Invitrogen). Fluorescence was detected using an inverted confocal microscope, Leica TCS SP8 (Leica). Quantification of detected signal per nucleus was performed using the CellProfiler Software (the Broad Institute of MIT and Harvard).

### Calcium flux measurement

Indo-1 solution was prepared by resuspending 50μg Indo-1 AM (Molecular Probes) in 25 μl DMSO (Sigma), 25 μl Pluronic-F127 (Invitrogen) and 113 μl FBS (Sigma). 2-5×10^6^ cells were incubated in 15 μl Indo-1 solution in 1ml IMDM (Gibco) + 1%FBS (Sigma) at 37°C for 45 min. Cells were washed and used at a concentration of 10^5^ cells in 500 μl RPMI (Thermo Fisher Scientific) + 1% FBS (Biochrom AG) and 1 mM CaCl_2_. 0.5 mM EGTA was added for the measurement. Basal calcium levels were acquired for 1min. Subsequently, cells were stimulated with 10ug/ml AffiniPure F(ab’)_2_ Fragment Goat Anti-Mouse IgM (JacksonImmuno), 2 μM FCCP (Agilent) or 1 μM ionomycin (ChemCruz). Data were acquired on Fortessa cytometer (BD) at the MDC Flow cytometry facility.

### ER tracker

To stain the endoplasmic reticulum 10^6^ primary B cells were stained in 100 μl 0.25 μM ER-Tracker Green or ER-Tracker Red in HBSS (0.137 M NaCl, 5.4mM KCl, 0.25 mM Na_2_HPO_4_, 0.44 mM KH_2_PO_4_, 1.3 mM CaCl_2_, 1.0 mM MgSO_4_, 4.2 mM NaHCO_3_) for 30 min at 37°C and washed before measurement on Fortessa cytometer (BD).

### Oxygen consumption rate

To assess mitochondrial function in primary B cells and DG75 cells, MitoStress Test using Seahorse XFe96 metabolic flux analyzer (Agilent) was used according to manufacturer’s instructions with the following modifications: Seahorse XF Cell Culture Microplates were pre-treated with poly-D-lysine and 10^6^ primary B cells or 10^5^ DG75 cells were plate in Seahorse XF Base Medium (Agilent) supplemented with 2mM L-glutamine (Agilent), 1mM sodium pyruvate (Agilent), for primary B cells 10mM glucose (Agilent), centrifuged at 200 x g at RT for 2 min without breaks and incubated for 30 min at 37°C in a CO_2_ –free incubator. 10 mM Glucose(Agilent) (in case of DG75), 1 μM Olygomycin (Agilent), 1 μM FCCP (Agilent), 0.5 μM antimycin A/rotenone (Agilent) were subsequently injected and oxygen consumption rate was measured. For primary B cells, optimized baseline reading and measurements were performed using 3 loops of mix (3 min), wait (0 min) and measure (3 min) (Traba *et al*, 2016).

Maximal oxygen consumption was calculated as OCR values obtained after FCCP treatment, spare respiratory capacity as the difference between OCR levels obtained after FCCP treatment and basal oxygen consumption, ATP production as the difference between basal OCR values and post-oligomycin treatment.

### Statistical analysis

Statistical analysis was performed using GraphPad Prism. The Shapiro-Wilk normality test was used to test for normal distribution. Tests used for analysis, number of animals and independent experiments are indicated in the respective figure legends. Experiments were considered independent from each other if cells were harvested and processed on separate days. For experiments performed with DG75 cells, bulk DG75 cells with acute IgH deletion were used and subsequently one IgH-KO clone was generated and used.

## Acknowledgements

The authors thank C. Salomon, G. Natale, J. Pempe and K. Petsch for excellent technical support; R. Lauhkonen-Seitz for administrative assistance; K. Rajewsky lab members for critical comments and suggestions, M. di Virgilio group for sharing reagents, MDC core technology platforms (Flow cytometry, Protein production) and MDC mouse facility for excellent support.

This research was supported by European Union’s Horizon 2020 research and innovation program under the Marie Sklodowska-Curie grant agreement No 752747 (to M.C.). J.J.’s research is supported by the German Research Foundation (DFG) project number: 419193696 and through the SFB1335. H.T is supported through the graduate school of the Max Planck Institute for Immunobiology and Epigenetics (IMPRS-IE) and through the SFB1335.

## Author contributions

M.C. designed, performed, analyzed most experiments and wrote the manuscript, H.T. performed PLA experiments, Y.Z. performed immunoblot experiments, A.Z. performed CRISPR-Cas9 mutagenesis on primary B cells, E.K. provided DG75 cell line and expertise, P.G. tested sgRNAs and K.S. developed Igβ targeting strategy. J.J. and K.R. supervised the project.

## Supplementary figure legend

**Supplementary Figure 1.**
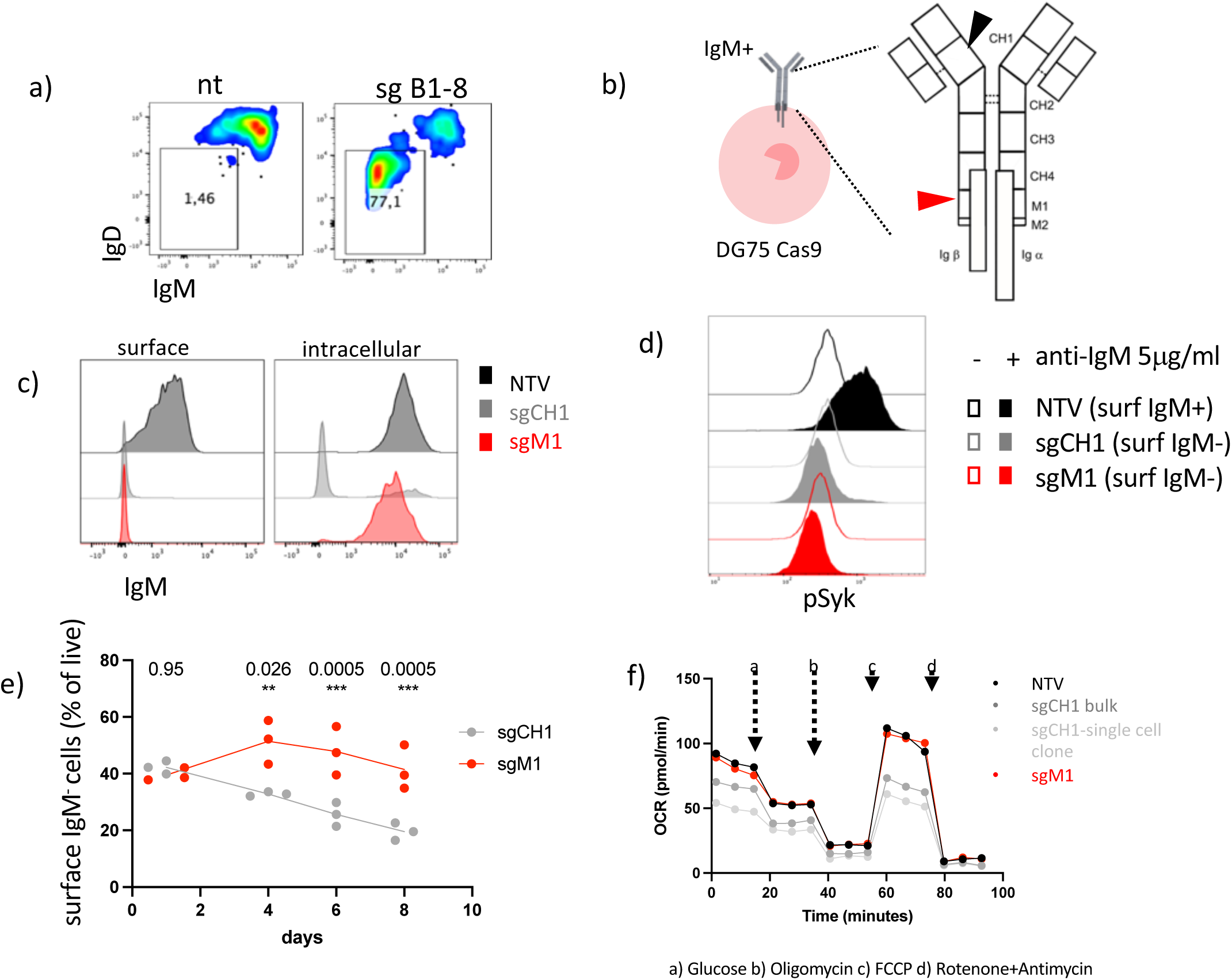
Intracellular Ig heavy chain provides competitive advantage to DG75 cells. (a) A representative flow cytometry analysis of mature resting B cells surface stained with anti-IgM and anti-IgD on d2 upon targeting using mock or sgB1-8; cells were cultured with BAFF (b) Schematic depiction of IgM targeting strategy in DG75 Cas9 cells with sgRNAs specific for CH1 and M1 region of Ig heavy chain. (c) A representative flow cytometry analysis of DG75 cells stained with anti-IgM on surface and intracellularly upon targeting with non-targeting sgRNA (NTV), sgCH1 or sgM1. (d) Phospho-flow analysis of pSyk in DG75 cells targeted with NTV, sgCH1 or sgM1 with or without anti-IgM stimulation. (e) Fitness of surface IgM-negative DG75 cells targeted with sgCH1 or sgM1. Results were tested for significance with ANOVA. (f) Oxygen consumption rates of DG75 cells targeted with NTV, sgCH1 (as bulk population or a single clone) or sgM1 measured with Mito stress test Seahorse assay. Data obtained from a minimum of 3 independent experiments.

**Supplementary Figure 2.**
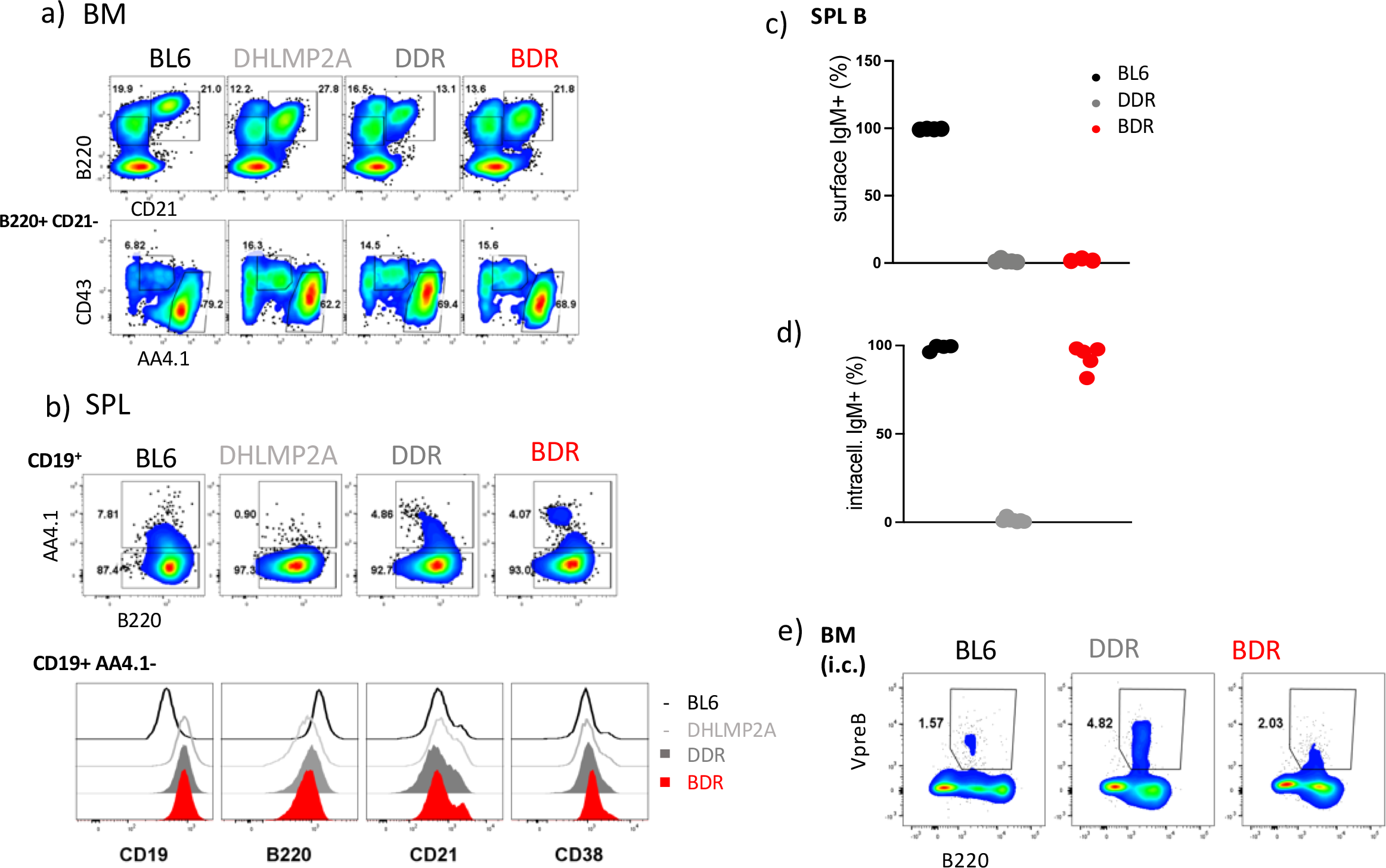
Mature splenic B cells in BDR mice have B2-like surface phenotype. (a) Flow cytometry analysis of BM cells from BL6, D_**H**_LMP2A, DDR and BDR mice stained with anti-B220, anti-CD21, anti-CD43 and anti-AA4.1 on surface. (b) Representative flow cytometry analysis of SPL B cells from BL6, D_**H**_LMP2A, DDR and BDR mice stained with anti-CD19, anti-AA4.1, anti-CD21, anti-CD38 and anti-B220 on surface. (c) Fraction of surface IgM^+^ B cells in spleen of BL6, DDR and BDR mice. (d) Fraction of intracellular IgM^+^ B cells in spleen of BL6, DDR and BDR mice. (e) Representative flow cytometry analysis of BM B cells from BL6, DDR and BDR mice stained with anti-CD179a (VpreB) intracellularly. Data obtained from a minimum of 3 mice in 3 independent experiments.

**Supplementary Figure 3.**
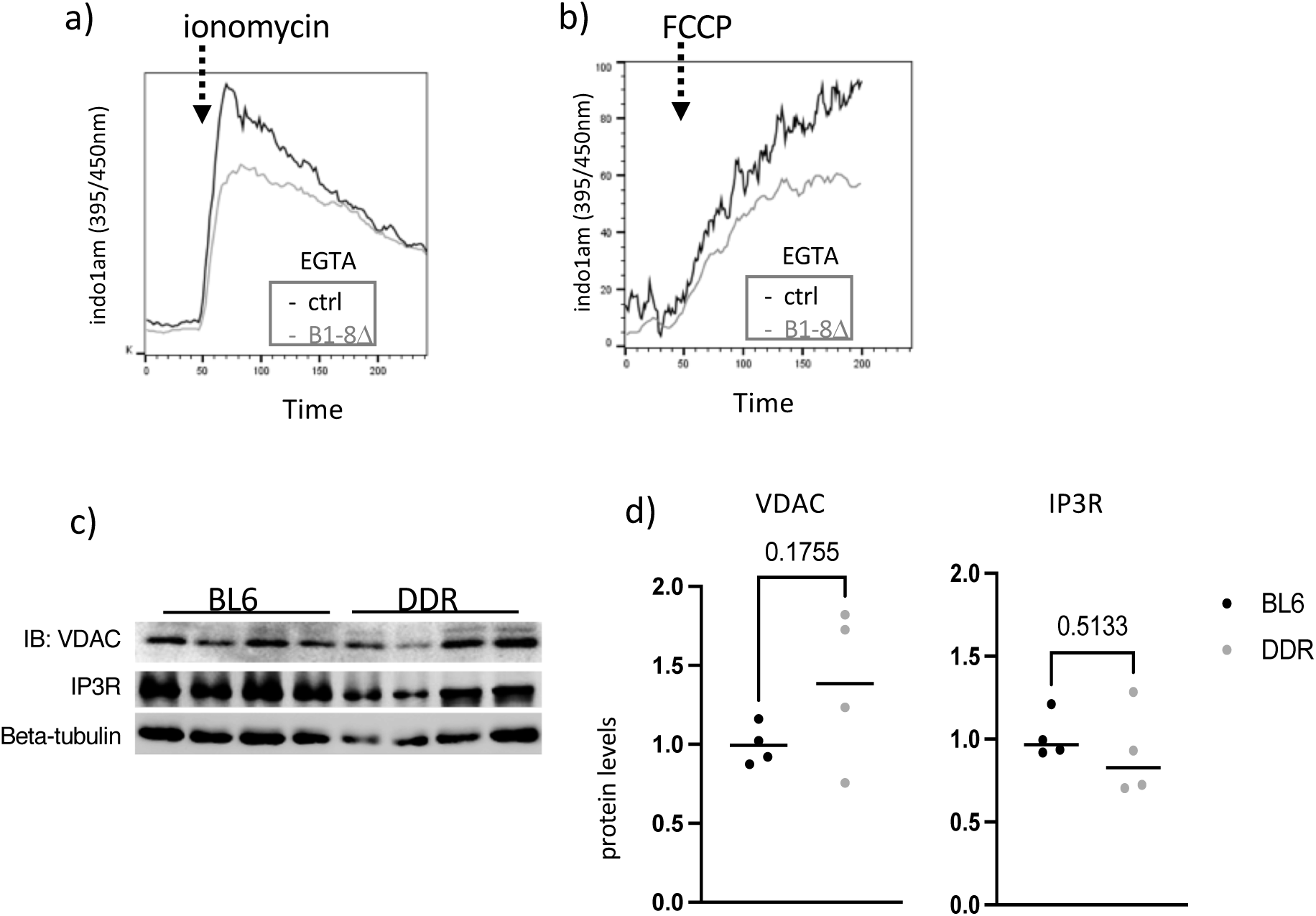
ER-resident Ig heavy chain improves ER homeostasis and MAMs. (a) Representative flow cytometry analysis of calcium efflux upon ionomycin stimulation in presence of EGTA in control (R26-hCD2^+^) and IgH-ko (B1-8Δ) B cells on d2 upon TAT-cre transduction. (c) Representative flow cytometry analysis of calcium efflux upon FCCP stimulation in presence of EGTA in control (R26-hCD2^+^) and IgH-ko (B1-8Δ) B cells on d2 upon TAT-cre transduction. (c) Immunoblot analysis of VDAC and IP3R protein in BL6 and DDR B cells. Beta-tubulin was used as loading control. (d) Unsaturated immunoblot images of VDAC and IP3R were quantified based on beta-tubulin from (c) using ImageJ and results were tested for significance with T-test. Data obtained from a minimum of 3 mice in 2 independent experiments.

